# Differential processing of delay versus uncertainty in male but not female 16p11.2 hemideletion mice

**DOI:** 10.1101/2023.10.04.560951

**Authors:** Gerardo R. Rojas, Abigail T. Heller, Nicola M. Grissom

## Abstract

Neurodevelopmental disorders are associated with differences in learning and motivation that can influence executive function, including behavioral flexibility and decision making. 16p11.2 hemideletion is a chromosomal copy number variant that is linked to neurodevelopmental disorders. 16p11.2 hemideletion in mice has been previously found to produce male-biased changes in reward learning, but the link between this and altered flexible decision making is poorly understood. We challenged 16p11.2 hemideletion mice with two reward-guided decision making tasks assessing flexible decision making under cost, delay and probability discounting. Both tasks elicited long-term changes in flexible decision making that separated 16p11.2 hemideletion males from wildtype males. In delay discounting,16p11.2 hemideletion males had a stronger, less flexible preference for the large reward at long delays, and this effect was reduced as wildtype males adjusted their preference to match that of the hemideletion males. In probability discounting, 16p11.2 hemideletion males initially had a similar preference for seeking improbable large rewards as did wildtype males, but over time began to prefer certainty to a greater extent than did wildtype males. Female mice discounted similarly for delayed or risky rewards regardless of the presence of the copy number variant. We have previously seen that male 16p11.2 hemideletion mice commit fewer nonreinforced responses than male wildtype mice in an operant setting, which we replicate here in delay discounting, while the introduction of risky rewards eliminates genotype differences in nonreinforced responses. Overall these data suggest that 16p11.2 hemideletion in males leads to differential processing of costs of delay versus inconsistency, with greater aversion to uncertainty than delays, and greater behavioral control by cues that consistently predict an outcome.

## 1. Introduction

Understanding how neurodevelopmental disorder-linked genes impact flexible decision making may shed light on the connections between fundamental neurobiology and diversity in cognition. One area of interest has been how neurodevelopmental disorders influence decision-making processes. It has been repeatedly observed that autism spectrum disorder symptoms may be partly explained by differential processing of environmental uncertainty (Lawson et al., 2017; Minassian et al., 2007; Sinha et al., 2014). Neurodevelopmental disorders are known to impact fundamental processes that support decision-making such as learning, motivation and attention (Dichter et al., 2012). Studies that have focused on the processing of rewards in ASD find reduced social motivation (Chevallier et al., 2012; Clements et al., 2018; Kohls et al., 2012), deficits in reward processing (Scott-Van Zeeland et al., 2010) and deficits in specific reward epochs like reward anticipation (Baumeister et al., 2023; Clements et al., 2018). In more complex situations which require probabilistic learning (e.g. Iowa Gambling Task), those with ASD show a broad trend of choices that resembles risk avoidance (South et al., 2014), but when closely examined could represent a difference in strategy (Zeif et al., 2023). This raises the question of whether applying different decision making tasks in the same individuals can reveal greater specificity in whether reward processing, costs, or uncertainty are most central in driving differences in flexible decision making.

Deletion of one copy of chromosomal region 16p11.2 in humans has been linked to diagnosis of autism and ADHD, and these individuals can exhibit language delays, social communication issues, and motor patterns typical of neurodevelopmental disorders regardless of whether or not specific diagnostic criteria are reached (Hanson et al., 2015; Rein & Yan, 2020; Walsh & Bracken, 2011). In mice, the 16p11.2 is highly conserved (Horev et al., 2011), and mice modeling this hemideletion show impacts in basal ganglia function (Portmann et al., 2014) and decreases NMDA receptor activity in the prefrontal cortex (Wang et al., 2018), both of which contribute to aspects of decision-making. Neurodevelopmental disorders such as autism spectrum disorder (ASD) and attention-deficit hyperactivity disorder (ADHD) are diagnosed at a higher rate in males than females (Loomes et al., 2017; Posserud et al., 2021). While some impacts of 16p11.2 hemideletion, including hyperactivity, are seen across sexes (Angelakos et al., 2017), sex-biased increases in several neurodevelopmental-disorder relevant domains, including sleep disturbances, anxiety-like behaviors, and reward learning alterations have been seen (Angelakos et al., 2017; Giovanniello et al., 2021; Grissom et al., 2018). Collectively, these data suggest that the impact of this copy number variant may be male-biased, but the extent to which this is true in flexible decision making is unknown.

Recently, we demonstrated mice are able to learn both delay and probability discounting (Rojas et al., 2022). Delay and probability discounting are valuable because they are complementary tasks that challenge animals with temporal or risky costs (Green & Myerson, 2004; McKerchar & Renda, 2012). In order to determine if 16p11.2 hemideletion mice are more sensitive to one type of cost, we tested mice on delay and probability discounting tasks. We had mice first undergo “Worsening” and then “Improving” versions of delay and probability discounting because the acclimation to an Improving schedule (i.e. going from no delay training to large delay testing) can obscure sensitivity to delay in models of neurodevelopmental disorders (Sjoberg et al., 2023).

Each schedule promotes a different choice pattern, which gives us the ability to compare whether delay or risk orientation alters choice in 16p11.2 hemideletion mice. We found that each task induced differences between 16p11.2 and wildtype males, but not females, but the differences depended on whether the task engaged the cost of delay or uncertainty. In delay discounting,16p11.2 hemideletion males had a stronger, less flexible preference for the large reward at long delays, and this effect was reduced as wildtype males adjusted their preference to match that of the hemideletion males. In contrast in probability discounting, 16p11.2 hemideletion males initially had a similar preference for seeking improbable large rewards as did wildtype males, but over time began to prefer certainty to a greater extent than did wildtype males. We have previously seen that male 16p11.2 hemideletion mice commit fewer nonreinforced responses than male wildtype mice in an operant setting, which we replicate here in delay discounting, while the introduction of risky rewards eliminates genotype differences in nonreinforced responses. Overall these data suggest that 16p11.2 hemideletion in males leads to differential processing of costs of delay versus inconsistency, with greater aversion to uncertainty than delays, and greater behavioral control by cues that consistently predict an outcome.

## 2. Methods

### 2.1 Subjects

Male 16p11.2 hemideletion mice (stock #013128) and female wildtype mice (stock #101043) were obtained from Jackson Laboratories and bred in order to generate mice of both genotypes for the experiments. Mice were housed in groups of 2-5 of mixed genotypes. Mice began experiments at approximately >120 days of age. Mice had free access to water and were food restricted with their home chow at 85–90% of their baseline weight. Mice were pre-exposed to the operant reinforcer, vanilla flavored Ensure, in their home cage for one day prior to training. Ensure was freely available to be licked from a bottle. Each group of mice were verified to have consumed a full bottle (148 ml) of Ensure. Behavioral testing took place Monday to Friday, and on Fridays, mice had free access to home chow. Animals were housed on a reverse light-dark cycle (9 am-11 pm) and were tested during the dark period. Animals were cared for in accordance with National Institute of Health guidelines and were approved by the University of Minnesota Institutional Animal Care and Use Committee.

### 2.2 Apparatus

We used 16 identical triangular touchscreen chambers, 8 per sex separated in different rooms for training and testing (Lafayette Instrument Co., Lafayette, IN). The touchscreen apparatus was located at the front of the chamber while liquid rewards were delivered at the back of the chamber. Schedules were administered and interactions with the screen were recorded via ABET-II software. All data were exported using ABET-II software. Touchscreens were limited by masks with five evenly spaced square holes. Ensure was diluted to 50% and delivered via peristaltic pump.

### 2.3 Touchscreen Training

Training was similar to our previous experiment (Rojas et al., 2022). Mice began with magazine training for 3 days and moved on to training schedules. For all training schedules, mice were only able to advance in trials if they performed the required action(s) designated by illumination by a 5-choice mask and collected the reward in the magazine at the back of the chamber. The touchscreen illuminated in the holes closest to the center where mice were required to nosepoke in order to advance the trial. Any responses on the outer two holes always resulted in no reinforcement at all stages. Reward delivery was accompanied by a light cue and the sound of the pump.

#### Center Only Fixed-Ratio 1

Mice experienced FR1 training for 40 days until a majority of them readily acquired a basic understanding to nosepoke for 7 μl of Ensure. Responses to the illuminated center hole resulted in reward delivery, all other responses were recorded but did not lead to reward delivery. Sessions ended after 30 minutes had elapsed.

#### Progressive Ratio

Mice intermittently experienced two sessions of progressive ratio. Mice were still required to selectively respond in the center hole for reinforcement. The first session occurred after 14 days of FR1 training. The second session occurred 37 days after FR1 training. Sessions advanced in an arithmetic sequence (1, 2, 4, 7, 11, 16, 21,…n). Each ratio had to be completed for three trials before advancing to the next in the sequence. Sessions terminated after animals failed to nosepoke for 5 minutes or after 60 minutes had elapsed. The last ratio completed became the animal’s breakpoint.

#### Chaining Center (Reinforced) to Left and Right

Animals began training to chain their response in order to build up to choices in discounting tasks. Sessions began with an illuminated center hole. Nosepoking the center hole resulted in 7 μl of Ensure, which upon collection illuminated the holes immediately to the left and right half of the center hole. Mice were then required to respond on either side for the same amount of Ensure on each side (28 μl). Training lasted for 24 days and sessions were 30 minutes in length.

#### Chaining Center (Unreinforced) to Left and Right

The next phase aimed to teach mice to respond in the center purely as an initiation response, resulting in no Ensure. The mice chose between left and right options which were still reinforced at the same volume. Mice experienced this schedule for 40 days and sessions were 40 minutes in length.

#### Chaining Center to Small and Large Magnitude Reward Options with ITI

The last stage of training before discounting was conducted before delay discounting in order to teach mice one option immediately left or right to the center hole resulted in a greater amount of Ensure (20 μl) than the other (5 μl). Additionally, the now large option resulted in a longer intertrial interval (ITI) of 30s. However, animals could choose no more than 2 of the same option in a row before they were forced to sample the other option. Mice learned these new concepts for 20 days and sessions ended after 60 minutes had elapsed or after mice cleared 60 trials.

### 2.4 Delay Discounting Phase

MIce underwent sequences of Worsening and Improving schedules as previously described (Rojas et al., 2022). One deviation from the previous study is that mice did not receive forced-choice trials in order to prioritize completion of delay blocks. Mice learned the Worsening schedule initially in which sessions began with a 0s delay for delivery Ensure after the large option, which temporally increased in cost as trials progressed (Figure 1). Sessions consisted of 40 free-choice trials divided into 5 blocks per delay (0, 4, 12, 20, and 28s delays). There was no delay to reinforcement for the small option, but mice did need to wait additional time to initiate the next trial (matched to the length of the delay trial). Mice proceeded to learn the Improving schedule in which sessions began with a 28s delay cost for Ensure after the large option, which temporally decreased in cost as trials progressed (28, 20, 12, 4, 0s delays). Small reward choices were again balanced to large option trial length by increasing the ITI length. Mice underwent 13 days of the Worsening schedule, 24 days of the Improving schedule, 10 days of a return to the Worsening schedule, and 10 days of a return to the Improving schedule.

**Figure 1.**
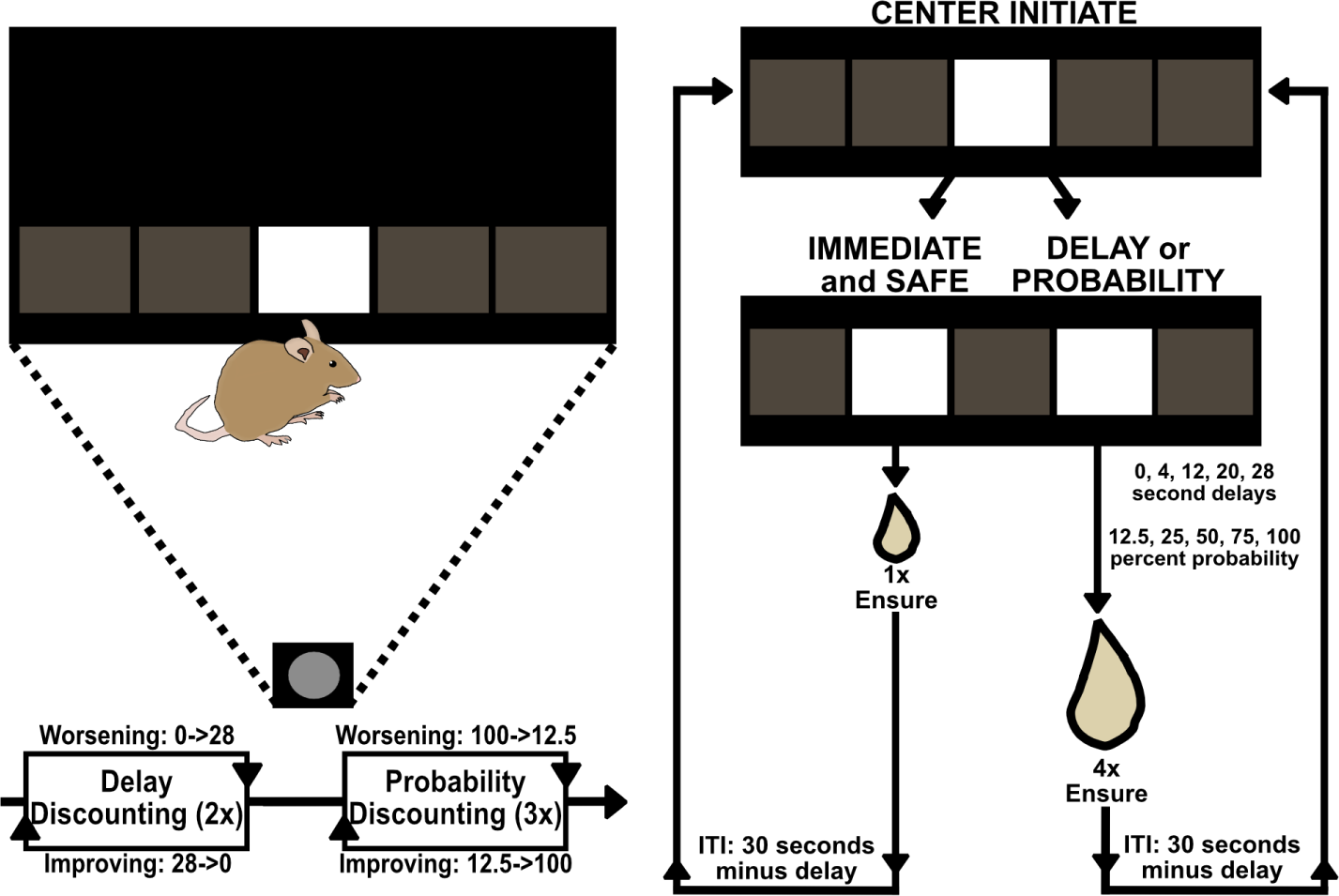
Sequencing of delay discounting and probability discounting. (Left-top) Mice were required to respond in illuminated areas of the touchscreen, limited to the 5 hole mask. The front of the chamber contained the touchscreen, the back of the chamber contained the magazine where Ensure was delivered. (Left-bottom) Mice experienced the Worsening schedule of delay discounting for 13 days before experiencing the Improving schedule for 24 days. They then repeated each schedule one more time before moving to probability discounting. Mice experienced the Worsening schedule for 10 days before experiencing the Improving schedule for 10 days. Mice then repeated each schedule two times before completing discounting. (Right) Sessions of discounting began with a center initiation nosepoke on a 5 hole mask. Mice were then presented with a small choice in which Ensure delivery was always safe and immediate (5 μl) and a large choice in which Ensure was 4x the magnitude (20 μl) but delivered in a delayed or probabilistic manner. ITIs were matched for trial length in delay and probability discounting. In delay discounting, ITIs were 30s minus the delay. In probability discounting, ITIs were 3s.

### 2.5 Probability Discounting Phase

Following delay discounting, mice then were exposed to the probability discounting task. Probability discounting training started with the Worsening schedule where 100% of trials resulted in delivery of the large reward, but uncertainty increased as trials continued (Figure 1). Sessions initially consisted of 40 trials for the first round of Worsening and Improving schedules, but switched to 80 trials in order to promote increased discounting of the reward. Sessions were divided into five probability blocks (100, 75, 50, 25, and 12.5% probability of large reward). Mice went on to learn the Improving schedule where odds of winning a large reward began at 12.5%, but probabilistically increased as trials progressed (12.5, 25, 50, 75, 100% probability of large reward). Mice underwent 10 days of the Worsening schedule, 10 days of the Improving schedule, 8 days of an initial return to the Worsening schedule, 15 days of an initial return to the Improving schedule, 10 days of a secondary return to the Worsening schedule, and 10 days of a secondary return to the Improving schedule. Data from the first round of probability discounting are excluded because of the decreased trial counts compared to the other rounds (40 vs 80) and because animals failed to show significant devaluation effects with increased risk.

## 3. Statistical Analysis

We constructed linear mixed models in order to analyze choice data and compare nonreinforced nosepokes. For analysis of large reward preference in delay discounting, we included fixed factors of Sex, Genotype, Schedule, Delay and their interactions (Sex x Genotype, Sex x Schedule, Sex x Delay, Genotype x Schedule, Genotype x Delay, Delay x Schedule, Sex x Genotype x Delay, Sex x Genotype x Schedule, Sex x Delay x Schedule, Genotype x Delay x Schedule, Sex x Genotype x Delay x Schedule). Random factors included a random slope of Schedule and random intercept of Subject. For analysis of large reward preference in probability discounting, we included fixed factors of Sex, Genotype, Schedule, Probability and their interactions (Sex x Genotype, Sex x Schedule, Sex x Probability, Genotype x Schedule, Genotype x Probability, Delay x Schedule, Sex x Genotype x Probability, Sex x Genotype x Schedule, Sex x Probability x Schedule, Genotype x Probability x Schedule, Sex x Genotype x Probability x Schedule). Random factors included a random slope of Schedule and random intercept of Subject. Comparisons of delay discounting and probability discounting preference within days were assessed using Welch’s t-tests (uncorrected for multiple comparisons). If mice were unable to reach at least 60% completion of trials on average across all days, those mice were excluded from analysis. Additional analysis of Worsening delay discounting within 28s delay of male mice was conducted with a linear mixed model consisting of main effects of Genotype and Training, and a Genotype x Training interaction term. There was also a random factor of Subject. Final group sizes for delay discounting analysis were as follows: 9 male wildtype mice, 9 male hemideletion mice, 10 female wildtype mice and 8 female hemideletion mice. Final group sizes for probability discounting analysis were as follows: 9 male wildtype mice, 9 male hemideletion mice, 10 female wildtype mice and 9 female hemideletion mice.

Analysis of nonreinforced touches in delay discounting and probability discounting included fixed factors of Sex, Genotype, Schedule and their interactions (Sex x Genotype, Sex x Schedule, Genotype x Schedule, Sex x Genotype x Schedule). There was a random intercept for Subject. Nonreinforced touches two standard deviations away from the mean were excluded from the analysis.

The R package *lmerTest* was used to fit linear mixed models. The Satterthwaite method was used to estimate degrees of freedom for omnibus F tests (main effects and interactions). Post-hoc comparisons were made using the *emmeans* R package with Tukey adjustment and Satterthwaite method for estimation of degrees of freedom. All graphs were produced using the *ggplot2* R package.

## 4. Results

### 4.1 Delay Discounting

#### Male 16p11.2 hemideletion mice resist devaluation effects of large delay

16p11.2 hemideletion mice in the past have been demonstrated to show hyperactivity and a delayed rate of reward learning (Angelakos et al., 2017; Grissom et al., 2018), both of which can correlate with a difference in reward valuation. 16p11.2 hemideletion mice have also been shown to stick to a choice rule once formed (e.g. perseverative choice in reversal learning; (Yang et al., 2015). In order to examine if this difference in learning extends to decision-making, we had 16p11.2 hemideletion mice experience both forms in order to identify possible vulnerabilities to delayed rewards. We present data from delay discounting progression to identify possible acquisition and mastery effects.

Across all of delay discounting, delay significantly reduced large reward preference (Figure 2A & 2B: main effect of Delay, p<0.001). Transitions from Worsening to Improving resulted in anchoring effect after initial training (Figure 2A & 2B (left): main effect of Schedule, F_(1, 36)_ = 14.894, p<0.001) and remained after additional training (Figure 2A & 2B (right): main effect of Schedule, F_(1, 36)_ = 11.773, p=0.002). Anchoring effects were prevalent at specific delays (Figure 2A & 2B: Schedule x Delay interaction, p<0.001).

**Figure 2.**
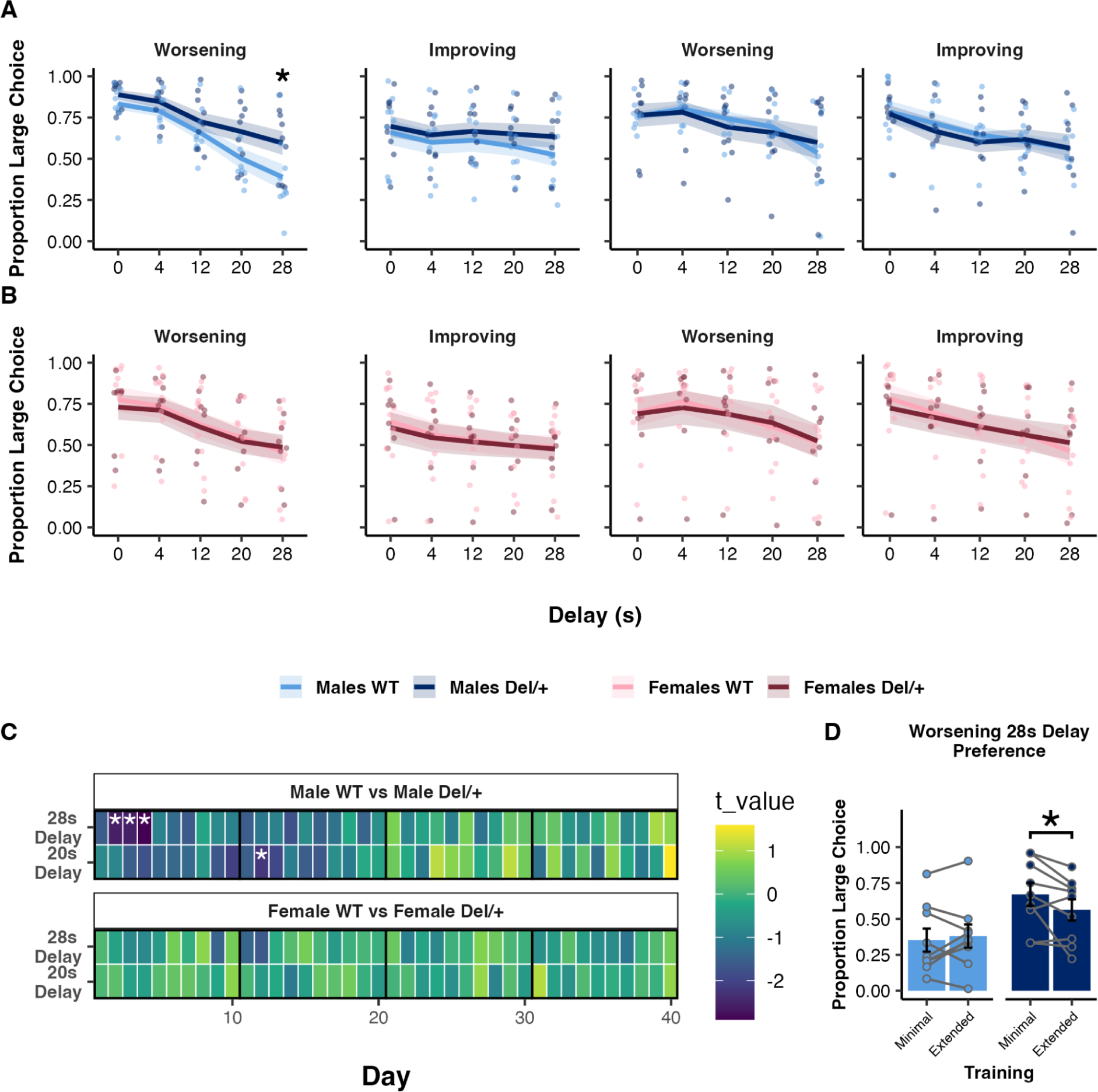
Male 16p11.2 hemideletion mice learned to shift reward preference with large delays. Mice transitioned from Worsening schedules (0s-> 28s delay) to Improving schedules (28s-> 0s delay) in order to assess schedule specific learning. (A) Male 16p11.2 hemideletion mice were relatively insensitive to longer delays compared to male wildtype mice (28s delay: p=0.0246). However, as male 16p11.2 hemideletion mice gained more experience with the task, they no longer exhibited delay specific differences compared to male wildtype mice. (B) Female mice exhibited similar discounting rates throughout all of delay discounting. (C) Discounting data is presented throughout days where the black vertical line indicates a change in schedule (Worsening to Improving, Improving to Worsening). When comparing learning history of those delays throughout training, it was evident that male 16p11.2 hemideletion mice preferred the large delayed option compared to male wildtype mice early but not late into training. In agreement with the overall trend with extended training, male genotype differences disappeared with additional training. D) Training data was split into a minimal timeframe (days 2-4) and an extended time frame (days > 4) to determine if wildtype mice became more like 16p11.2 hemideletion mice or vice versa within the first round of Worsening delay discounting. Our data confirms male 16p11.2 hemideletion mice but not wildtypes significantly alter their preference point (p=0.0215). Shaded areas in A and B and error bars represent standard error from the mean. Asterisks represent statistical significance of p<0.05.

Within the first round of Worsening and Improving delay discounting, 16p11.2 hemideletion mice showed evidence of resisting devaluation effects caused by delays to reward (Figure 2A & 2B (left): Genotype x Delay interaction, F_(4, 288)_ = 3.207, p=0.0134). Upon comparing genotype differences in discounting within sexes, we found male 16p11.2 hemideletion mice repeatedly chose large delayed rewards more often than male wildtype mice when the delay was largest (Figure 2A (left): male 20s delay, p=0.0721 NS; male 28s delay, p=0.0237). Although evidence suggests there might be sex differences in delay discounting (Figure 2A & 2B (left): Sex x Delay x Schedule interaction, F_(4, 288)_ = 2.589, p=0.0370), post-hoc comparisons do not support within schedule differences.

In order to determine the strength of this male-specific difference in delay discounting, we compared preference rates across multiple days. Male 16p11.2 hemideletion mice chose the large reward more often than male wildtype mice specifically early into training, but adjusted their discounting to become as sensitive to delays as wildtype mice (Figure 2C: 28s delay: day 2, t_(13.604)_ = −2.576, p=0.022; day 3, t_(15.585)_ = −2.565, p=0.021; day 4, t_(15.197)_ = −2.898, p=0.011). These data indicate male-specific differences in discounting are minimized as mice gain more experience on the task. This is in stark contrast to female mice who consistently chose the large reward at a similar opportunity cost ratio (Figure 2C: p>0.05). In order to determine if the difference in discounting at the 28s delay within the first round of Worsening discounting was due to a change in the male wildtypes or male 16p11.2 hemideletion mice, we compared their training history and grouped them into a Minimal phase (days 2-4) and an Extended phase (days > 4). We discovered that it was male 16p11.2 hemideletion mice that significantly shifted their preference over the course of the first round of Worsening delay discounting (Figure 2D: Genotype x Training interaction, F_(1, 18)_ = 5.027, p=0.0378). Post-hoc comparisons confirm this was only significant within male 16p11.2 hemideletion mice (Figure 2D: p=0.0215)

#### Male 16p11.2 hemideletion mice commit fewer trial initiation errors, indicative of action inhibition through a nonreinforcement period

Male 16p11.2 hemideletion mice have previously been shown to use different response strategies to attain rewards in that they tend to make significantly less nonreinforced responses in a simple continuous reinforcement task (Grissom et al., 2018). As such, we measured those nonreinforced touches during different epochs of the task in order to determine if nonreinforced touches were differently influenced between genotypes as a function of task progression. We present nonreinforced responses during the choice period throughout the delay discounting session as a measure of initiation perseveration.

Consistent with previous research, 16p11.2 hemideletion mice made significantly fewer nonreinforced nosepokes in the center (initiation) hole during the choice period for both the first half of discounting (Figure 3A (left half of males and females): main effect of Genotype, F_(1, 32)_ = 5.711, p=0.0227) and second half of discounting (Figure 3 (right half of males and females): main effect of Genotype, F_(1, 34.742)_ = 8.010, p=0.0077). Nonreinfored nosepokes were initially increased on the Improving Schedule compared to the Worsening Schedule (Figure 3 (left half of males and females): main effect of Schedule, F_(1, 33)_ = 4.263, p=0.0469) but that difference diminished with additional experience. Post-hoc tests revealed this difference was mainly due to more nonreinforced touches made by female 16p11.2 hemideletion mice on the Improving Schedule (Figure 3 (left half of females), p=0.0178). Male 16p11.2 hemideletion mice made less nonreinforced nosepokes for both the first round of discounting (Figure 3 (left half of males): Worsening, p=0.0514 NS; Improving, p=0.0441) and the second round of discounting (Figure 3 (right half of males): Worsening, p=0.0015; Improving, p=0.0219). Females exhibited no such genotype differences (p>0.05).

**Figure 3.**
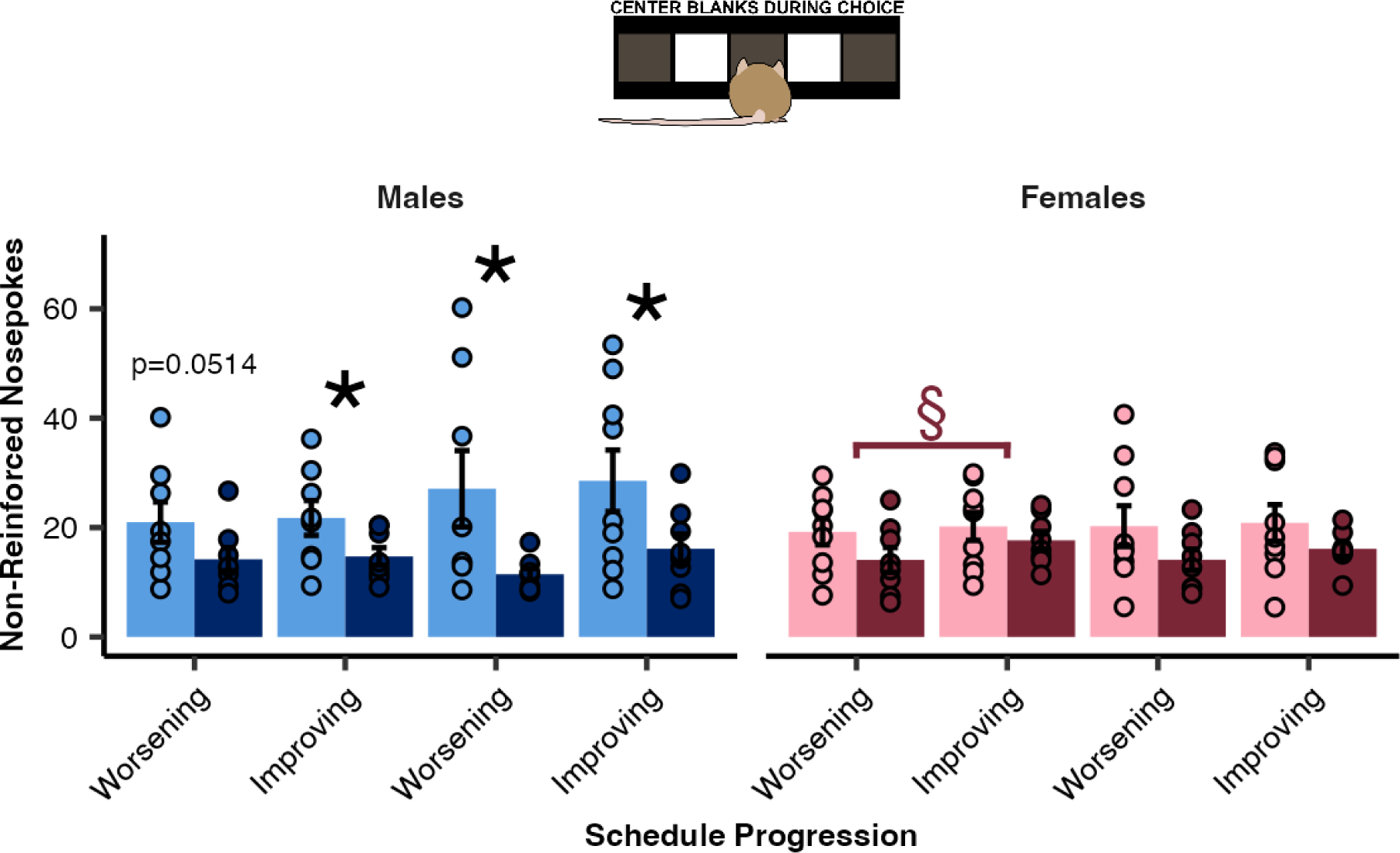
Male 16p11.2 hemideletion mice withheld center nosepokes during the choice epoch of delay discounting. For all of delay discounting, male 16p11.2 hemideletion mice make less initiation errors than male wildtype mice. Female 16p11.2 hemideletion mice make more nonreinforced nosepokes when the schedule starts with a large delay and progressively becomes shorter (Improving). Error bars represent standard error from the mean. An asterisk represents a genotype effect of p<0.05, a double s represents a schedule effect within a genotype (indicated by color) of p<0.05.

### 4.2 Probability Discounting

#### Male genotype differences in risk preference arise with experience of large risk

Reward uncertainty is another form of cost that can modulate the value of a reward. A prominent theory within the study of neurodevelopmental disorders implicates the importance of stable cues in learning environments (Sinha et al., 2014). We challenged 16p11.2 hemideletion mice with probability discounting in order to determine if they can adjust to the uncertainty of reward delivery coming off of a deterministic task like delay discounting. We again present discounting data to emphasize the progression of learning.

For all probability discounting sessions, we found an expected significant main effect of probability indicating a tendency to reduce large reward preference as risk increased (Figure 4A & 4B: main effect of Probability, p<0.001). Anchoring effects were clearly present within the first round of probability discounting (Figure 4A & 4B (left): main effect of Schedule, F_(1, 37)_ = 6.663, p=0.0139), but the influence of anchoring diminished over additional training (Figure 4A & 4B (right): p>0.05). However, for both rounds there was clear evidence of probability specific differences in discounting between schedules (Figure 4A & 4B: Schedule x Probability interaction, p<0.001). Male mice generally endured risk more often for large rewards than female mice for the first round of probability discounting (Figure 4A & 4B (left): main effect of Sex, F_(1, 37)_ = 4.571, p=0.0392), a trend which was present but not significant with additional training (Figure 4A & 4B (right): main effect of Sex, F_(1, 37)_ = 3.942, p=0.0546). These increased male risky decisions were prominent at multiple probabilities (Figure 4A & 4B: Sex x Probability interaction, p<0.001).

**Figure 4.**
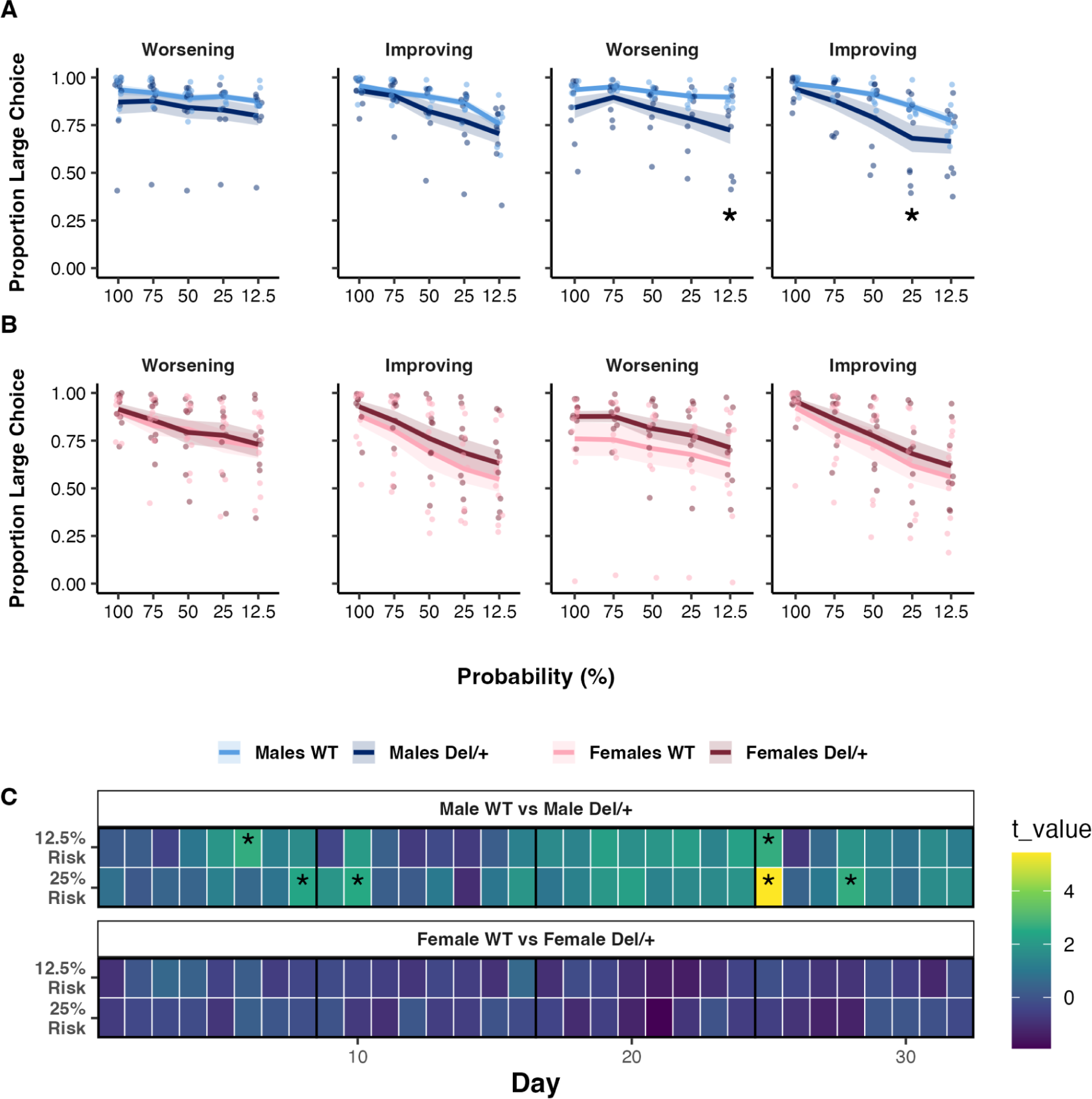
Male 16p11.2 hemideletion mice became slightly more risk avoidant with probability discounting training. Mice transitioned from Worsening schedules (100%-> 12.5% probability) to Improving schedules (12.5% probability-> 100% probability) in order to assess schedule specific learning. (A) Mice show greater cost endurance for large uncertain rewards compared to delayed large delayed rewards. 16p11.2 hemideletion and wildtype mice discount similarly initially on both Worsening and Improving schedules. (B) However, male 16p11.2 hemideletion mice become steeper than male wildtype mice as risk increases. Specifically, male 16p11.2 hemideletion mice prefer smaller rewards more often than male wildtype mice when risk of reward delivery is highest (12.5%: p=0.0354). (C) Discounting data is presented throughout days where the black vertical line indicates a change in schedule (Worsening to Improving, Improving to Worsening). Training history analysis does hint at some transient increased risk sensitivity that becomes more apparent with additional training. Shaded areas in A and B and error bars represent standard error from the mean. Asterisks represent statistical significance of p<0.05.

Male mice displayed differences in risk processing in the second half of probability discounting. Male 16p11.2 hemideletion mice were more risk averse when uncertainty of reward delivery was high on the Worsening Schedule (Figure 4A (right): male 12.5% Probability, p=0.0369) and the Improving Schedule (Figure 4A (right): male 25% Probability, p=0.0274).

We compared discounting preferences across days in order to assess the consistency of these effects. Differences do arise early and late into training of probability discounting when risk is greatest (Figure 4C: 12.5%: day 6, t_(8.799)_ = 2.649, p=0.027; day 25, t_(12.359)_ = 2.657, p=0.020). Interestingly, differences were more apparent when the expected value of the large risky option was in conflict with the value of the small option (Figure 4C: 25%: day 8, t_(8.131)_ = 2.407, p=0.042; day 10, t_(8.814)_ = 2.524, p=0.033; day 25, t_(9.843)_ = 5.435, p<0.001; day 28, t_(10.751)_ = 2.649, p=0.023).

#### Uncertainty of reward delivery erases male genotype differences in nonreinforced responding

Delay discounting adds a delay to reinforcement but is ultimately still a deterministic task. Animals tend to use different strategies when delivery of reinforcement is uncertain. Therefore we measured nonreinforced nosepokes through all epochs of probability discounting to determine if the previously measured genotype difference is still present. We present nonreinforced responses during the choice period throughout the probability discounting session as a measure of initiation perseveration and to compare to delay discounting.

Male genotype differences in nonreinforced nospokes committed in the center hole during the choice period were not significant, but were still present early into probability discounting (Figure 5 (left half of males): Worsening, p=0.0997 NS; Improving, p=0.0818 NS) and became less apparent as mice gained additional experience. While females were still similar in their pattern of nonreinforced nosepokes, female wildtypes adjusted to the uncertainty of the task by the end of training by reducing their nonreinforced nospokes (Figure 5 (right half of females): p=0.0284). Comparing these findings to the delay discounting nonreinforced nosepokes suggests task uncertainty disrupts or influences male 16p11.2 hemideletion specific action patterns.

**Figure 5.**
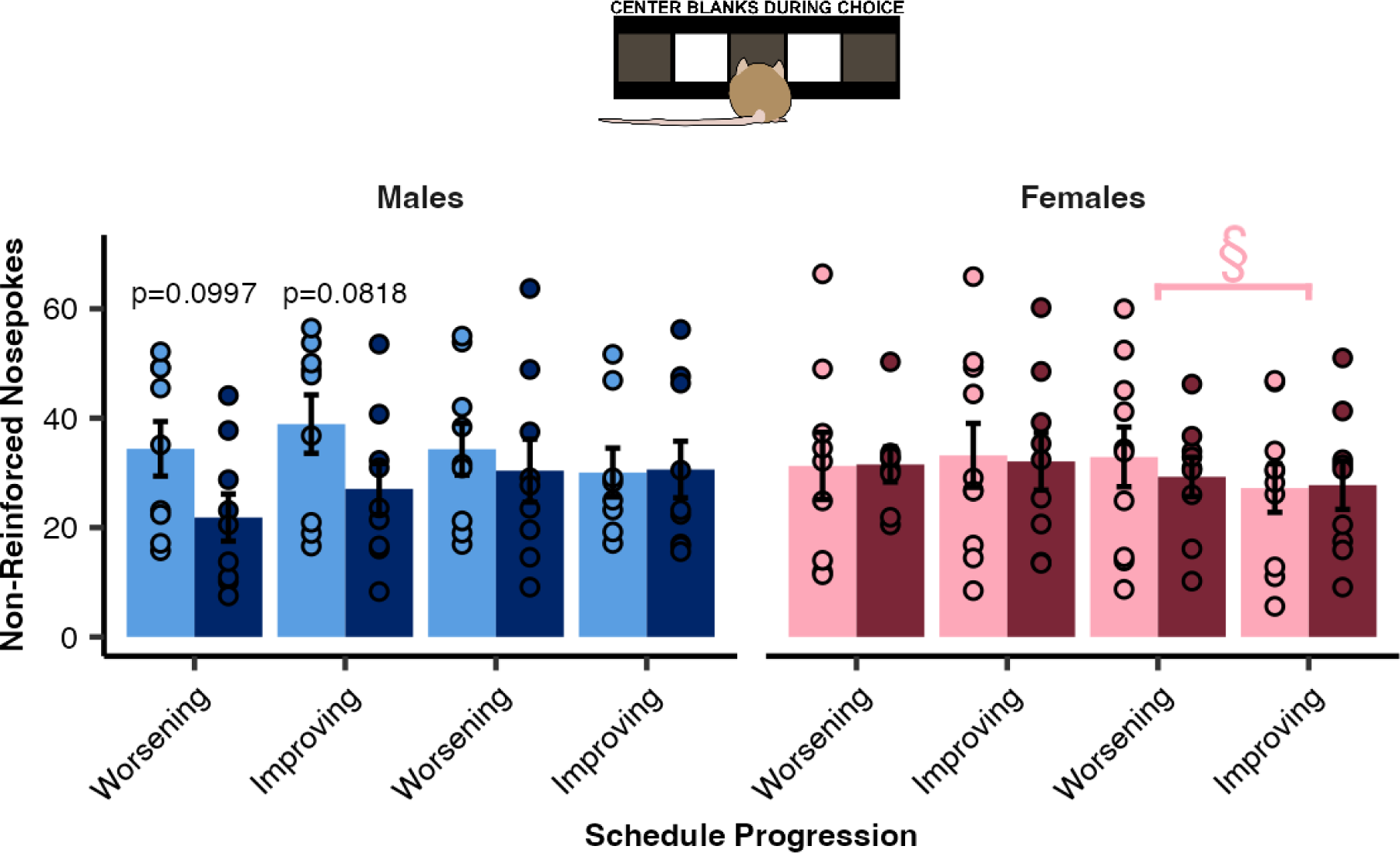
Male 16p11.2 hemideletion initiation mistakes increased upon training on probability discounting. Upon switching from delay discounting to probability discounting, male 16p11.2 hemideletion mice committed just as many initiation errors as male wildtype mice. Female wildtype mice adapt to the uncertainty of the task at the end of probability discounting by reducing the amount of initiation errors they commit. Error bars represent standard error from the mean. Double s represents a schedule effect within a genotype (indicated by color) of p<0.05.

## 5. Discussion

At the beginning of our study, we set out to challenge decision-making in 16p11.2 hemideletion mice in order to determine the impact of a copy number variant related to neurodevelopmental disorders. We chose delay and probability discounting tasks because they assess different aspects of reward-guided decision-making, of which is altered in neurodevelopmental disorders (Clements et al., 2022; Damiano et al., 2012; Mosner et al., 2017; Nissan et al., 2023). Using our battery of decision-making tasks, we provide evidence that 16p11.2 hemideletion impacts choices in two types of discounting tasks. Previous research found male 16p11.2 hemideletion mice are slower to form action-outcome relationships, resulting in delayed instrumental learning in a simple reinforcement task (Grissom et al., 2018). Our results corroborate those findings and expand upon them in a choice paradigm where task demands progressively shift within a session. We found that male 16p11.2 hemideletion mice exhibited resistance to temporal costs and delayed sensitivity to probabilistic costs compared to male wildtype mice. Male genotype differences in response to temporal delays occur initially with minimal experience, while the effects of reward uncertainty emerge with extensive experience. Patterns of nonreinforced responses during the choice period shift between tasks where male 16p11.2 hemideletion mice noticeably commit less trial initiation errors. Our data suggests male 16p11.2 hemideletion mice engage with their environment in different ways than do male wildtype mice and display temporarily enduring patterns of choice.

Our results are in agreement with previous findings on 16p11.2 hemideletion mice emphasizing slow learning and response inflexibility. Previous studies report delayed operant learning of which we find a parallel to here with male 16p11.2 hemideletion mice taking longer than male wildtype mice to shift their choice in response to large delays (Grissom et al., 2018). We found evidence of initially rigid choice similar to previous studies emphasizing inflexibility and perseverative choice in 16p11.2 hemideletion mice (Yang et al., 2015). These papers suggest a general challenge with adapting to reward-guided tasks, but do not elucidate why. Slower or different adaptation to costs is a phenotype observed in these mice and in neurodevelopmental disorders generally (Mussey et al., 2015; Yechiam et al., 2010; Zeif et al., 2023). Some research suggests increased sensitivity to losses can lead to a difference in choice in probabilistic tasks (Gosling & Moutier, 2018). Uncertainty avoidance can form when predictions are violated, resulting in what appears to be increased sampling or a tendency to shift towards certainty (Bervoets et al., 2021; Sinha et al., 2014; Zeif et al., 2023). Similarly, when reward certainty decreases in probability discounting, male 16p11.2 hemideletion mice significantly shift their choices toward the small certain choice. These results suggest exploratory behavior and decisions in 16p11.2 hemideletion mice could be modulated by uncertainty or risk of reward, which might also make it difficult to acquire new responses. Taken together, our results suggest male 16p11.2 hemideletion mice assess reward value in a manner that is different from wildtype mice and that is experience and uncertainty dependent. Future research could examine decision-making in 16p11.2 hemideletion mice in a dynamic setting such as in bandit tasks (Chen et al., 2020, 2021). This will enable researchers to determine the stability of 16p11.2 hemideletion-induced strategies and periods of possible perseveration as reward contingencies shift.

Next, we decided to measure nonreinforced touches throughout discounting in order to examine if exploratory behavior or approaches to learning about choice contingencies differed between 16p11.2 hemideletion and wildtype mice. One feature of operant behavior in 16p11.2 hemideletion that we observed here is a differential rate of nonreinforced responding, where hemideletion males make fewer nonreinforced responses than do males. Grissom et al. (2018) previously saw a similar pattern in a different task (5-choice serial reaction time task) and different operant chamber format (9-hole nosepokes versus touchscreen). This suggests that one source of male-biased impacts of 16p11.2 hemideletion is a reduction of unnecessary actions that wildtype males exhibit. Two possible explanations for nonreinforced responding in general that may differ across genotypes are differences in motivation/hyperactivity, or differences in attentional enhancement. To address the first hypothesis, anticipatory responses leading up to choice can be thought to reflect a type of general “impulsivity” (Dalley et al., 2011; Hogarth et al., 2012), while reduced motivation and vigor of actions are elicited when rewards are less frequent or more delayed (Ko & Wanat, 2016; Mohebi et al., 2019; Nicola, 2010). Through this lens, male 16p11.2 hemideletion mice may exhibit less effort towards unnecessary responses when rewards are temporally distal (in delay discounting) compared to when they are more immediate and uncertain (in probability discounting). Prior work has shown reduced responding in 16p11.2 hemideletion male mice in a progressive ratio task, consistent with this hypothesis (Grissom et al., 2018). The second hypothesis is that 16p11.2 hemideletion males may have greater attentional control and/or ability to inhibit prepotent responses. (Openshaw et al., 2023) recently used a continuous performance task and found that male 16p11.2 hemideletion mice performed with greater accuracy as measured by hit rate (i.e. responding during a correct stimulus) compared to male wildtype mice. Because the nonreinforced responding we measured occurred after the center hole was no longer illuminated, it may be that wildtype males have their responding under less stringent control of the cue than do 16p11.2 hemideletion males. One strategy for rational agents in discounting tasks is to decrease attention during nonrewarding periods and increase attention when the reinforcer becomes available again (Mikhael et al., 2021). For male wildtypes, extra responses may be one way in which to combat the ambiguity of the ITI period. Male 16p11.2 hemideletion mice rely more on the presence of response cues that signal for trial phases. However, when the reliability of reinforcement decreases such as in probability discounting, male 16p11.2 hemideletion mice start to adopt the same behavioral pattern as male wildtype mice. Future task designs may wish to explicitly test whether changes in motivation or changes in attentional or cue-regulated control form the greater contribution of behavioral differences in 16p11.2 hemideletion males.

Our work supports growing evidence that 16p11.2 hemideletion impacts males more than females especially in reward-related scenarios. Males show genotype differences discounting while females are generally unaffected in their choice behavior across both tasks similar to other types of reinforcement learning settings (Grissom et al., 2018). Contrary to findings of increased hyperactivity in 16p11.2 hemideletion mice, we found evidence of increased behavioral control (Angelakos et al., 2017). However as the authors point out themselves, 16p11.2 hemideletion-induced hyperactivity is context dependent. General locomotion in small operant chambers tend to be similar regardless of genotype (Grissom et al., 2018). Instead, 16p11.2 hemideletion mice committed fewer nonreinforced nosepokes similar to previous findings (Grissom et al., 2018). One group found evidence of anxiety-like behavioral and neurological responses after a fear-inducing event in female 16p11.2 hemideletion mice (Giovanniello et al., 2021). Probability discounting is one task in which risk attitudes affect discounting rates (Shead & Hodgins, 2009), so we expected female genotype differences to potentially be exposed there. However, contrary to those expectations female mice discounted at similar rates indicating the need for more overt punishments for exposing possible genotype differences. These results highlight how this CNV is affected by background, genotype and sex (Grissom et al., 2018; Horev et al., 2011; Portmann et al., 2014). Given that 16p11.2 hemideletion in humans has different impacts based on other genotypes and across genders (Chawner et al., 2019; Duyzend & Eichler, 2015; Hanson et al., 2015), this highlights the importance of understanding cognitive and neural impact of neuropsychiatric-disorder linked copy number variants as a function of individual differences.

## Acknowledgements

We would like to thank Ted Abel for his helpful comments and guidance for this project and Evan Knep for assisting with the completion of the experiments.

